# Comparison of the 3D-IR - BTFE method and the conventional method in the head MRI contrast

**DOI:** 10.1101/211763

**Authors:** Daisuke Hirahara

## Abstract

MRI using gadolinium contrast media is useful in diagnosis; however, nephrogenic systemic fibrosis is a serious side effect of gadolinium exposure. Moreover, it turns out that gadolinium deposits in the brain. This has escalated the necessity for a suitable method to use gadolinium contrast media. I developed a new imaging method that had excellent contrast. This study examined the usefulness of that new imaging method and found the method is highly effective.

## 1) Background

Exact diagnosis of a metastatic brain tumor is important for stage judging and future treatment policy. The first choice for metastatic brain tumor imaging is MRI, which uses double the amount of gadolinium contrast media.^1–4^ MRI using gadolinium contrast media plays an important role in the differential diagnosis of other brain tumors. However, it has been reported that nephrogenic systemic fibrosis (NSF) is a serious side effect of gadolinium contrast media. One cause of NSF is that contrast media chemical structure is not a macro ring contexture. Another cause is the amount used. Therefore, a renal function check is now required. Moreover, the amount used and the interval of contrast media were optimized. Kanda et al. 2013 showed that with a contrast media that is not a macro ring contexture, the risk to a brain is clear, and did not recommend the use of any contrast media other than those with a macro ring contexture. Subsequent research revealed that macro ring contexture agents also deposited slightly in the brain. To avoid risks, it is necessary to use gadolinium contrast media appropriately.^5–7^ After gadolinium imaging, a T1 image of the spin echo method was used. The three-dimensional gradient echo imaging method (3D-GRE) T1 image was also used. The MPRAGE method was used in 3T by the influence of SAR and T1 extension.^8^

## 2) Purpose

With an imaging method that emphasizes the contrast effect more than the conventional imaging method, there is a high possibility of reducing the risk of side effects due to the reduction of the gadolinium contrast agent. Further, there is a high possibility of improving the lesion detection rate due to the increase in contrast. In this study, 3D-IR (IR) and the inversion recovery method (IR) were combined to obtain a T1 emphasis based on a balanced turbo field echo (BTFE) sequence, which is a coherent gradient echo method, which is a high signal-to-BTFE imaging method was prepared and the conventional imaging method and contrast were examined.^9^

## 3) Apparatus and method

This survey was conducted in accordance with the Declaration of Helsinki; subjects orally consented. There are eight examples of metastatic brain tumor searches. The equipment was a superconducting 1.5 Tesla MRI machine (Intera Achieva Nova, PHILIPS) using an 8CH SENSE head coil. The contrast media used was gadolinium (HP-DO 3A) (0.4 ml/kg) for the patient. EZR was used for statistical processing.^10^ I used a slice thickness of 3 mm of T1 emphasis picture (SE-T1WI) for the two-dimensional spin echo method and a slice thickness of 1 mm for the three-dimensional gradient echo method (3D-GRE). 3D-IR-BTFE also used a slice thickness of 1 mm. I reconstructed in a slice thickness of 3 mm so that SE-T1WI could be compared with an imaging method with a slice thickness of 1 mm. To eliminate the order effect, the order of each imaging method after gadolinium imaging was random. Considering the distribution of gadolinium contrast media in the brain, I started the image pick-up 5 minutes after pouring the contrast media. The main imaging conditions are shown in Tables 1 and 2.

**Table 1:** shows the main imaging parameters of the SE-T1 WI.

|  |  |  |  |
| --- | --- | --- | --- |
| FOV | 230 | Slice thickness | 3mm |
| RFOV | 80 | Slice gap | 0mm |
| MATRIX | 256 | Scan mode | MS |
| RECON | 512 | Technique | SE |
| Scan% | 70 | TR | 462 |
| SENSE | No | TE | 15 |
| Slices | 48 | Flip angle | 90 |
| NSA | 2 | Scan time | 6min49sec |

**Table 2:** shows the main imaging parameters of the created 3D-IR-BTFE.

|  |  |  |  |
| --- | --- | --- | --- |
| FOV | 230 | Slice thickness | 1mm |
| RFOV | 80 | Slice gap | 0mm |
| MATRIX | 224 | Scan mode | 3D(IR delay:1200) |
| RECON | 256 | Technique | FFE(TFE factor:256) |
| Scan% | 110 | TR | 4.3 |
| SENSE | YES (P:2.5,S:1.0) | TE | 2.2 |
| Slices | 140 | Flip angle | 60 |
| NSA | 2 | Scan time | 3min38sec |

This evaluation compared contrast. Contrast comparison set the area of interest (ROI) as a pathological change in brain substance and evaluated it.

## 4) Result

Part of an obtained image is shown in Fig. 1.

**FIG. 1:**
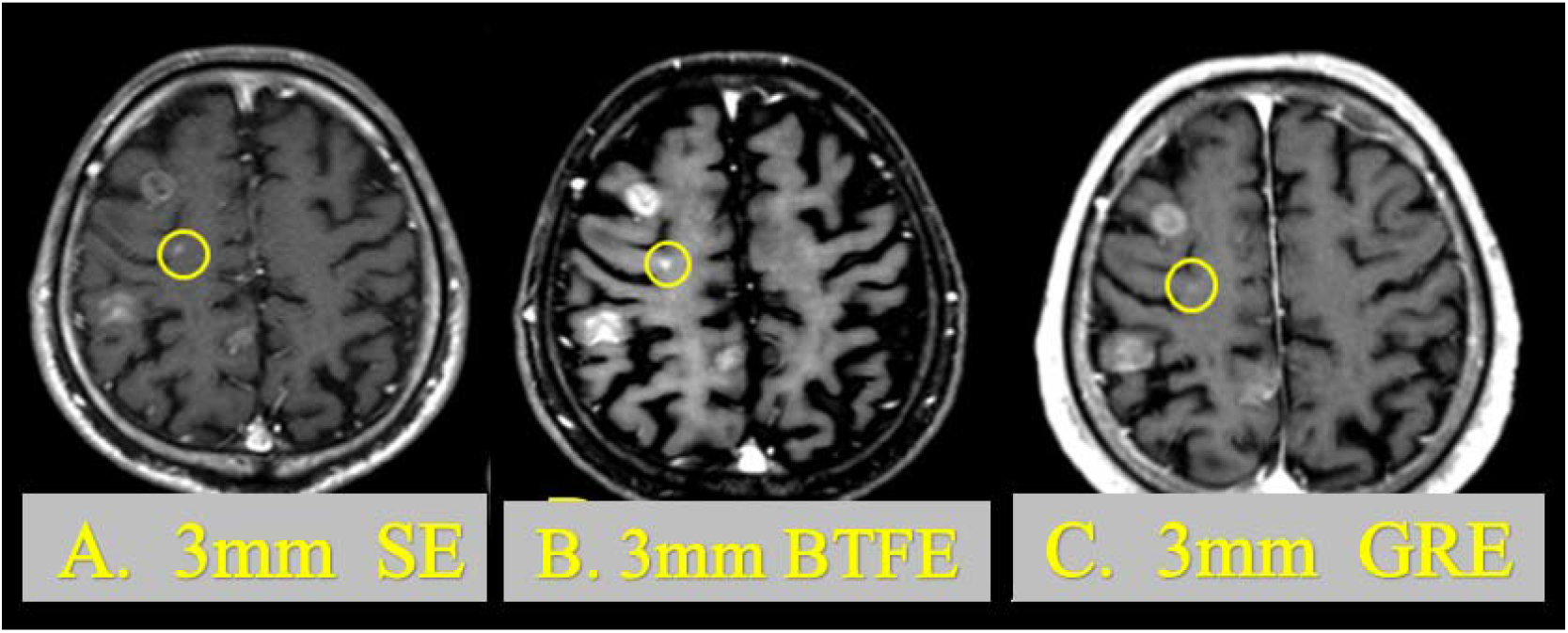
shows a photographed image. A is SE-T1 WI of 3 mm after imaging. B is 3D-IR-BTFE of 3 mm after imaging. C is 3D-IR-BTFE of 1 mm after imaging.

An example of a setting a ROI is shown in Fig. 2.

**FIG. 2:**
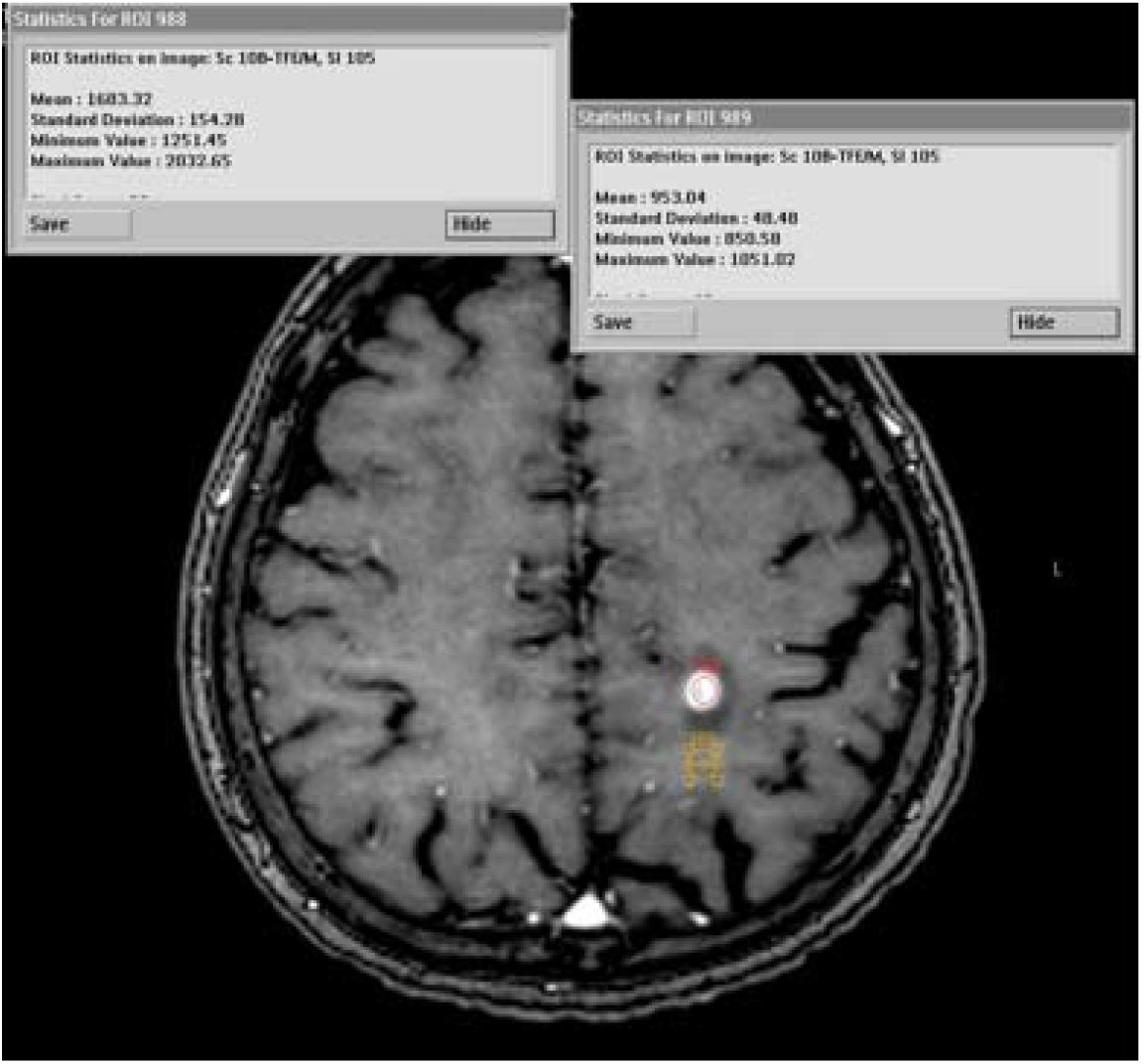
shows a method for setting the actual region of interest.

The difference in the average values of SE-T1WI and 3D-IR-BTFE is shown in Fig. 3. The difference test of the average value was t (19) = - 8.252, p < .01 (p = 0.0000001), d = - 9.64.

**Fig. 3:**
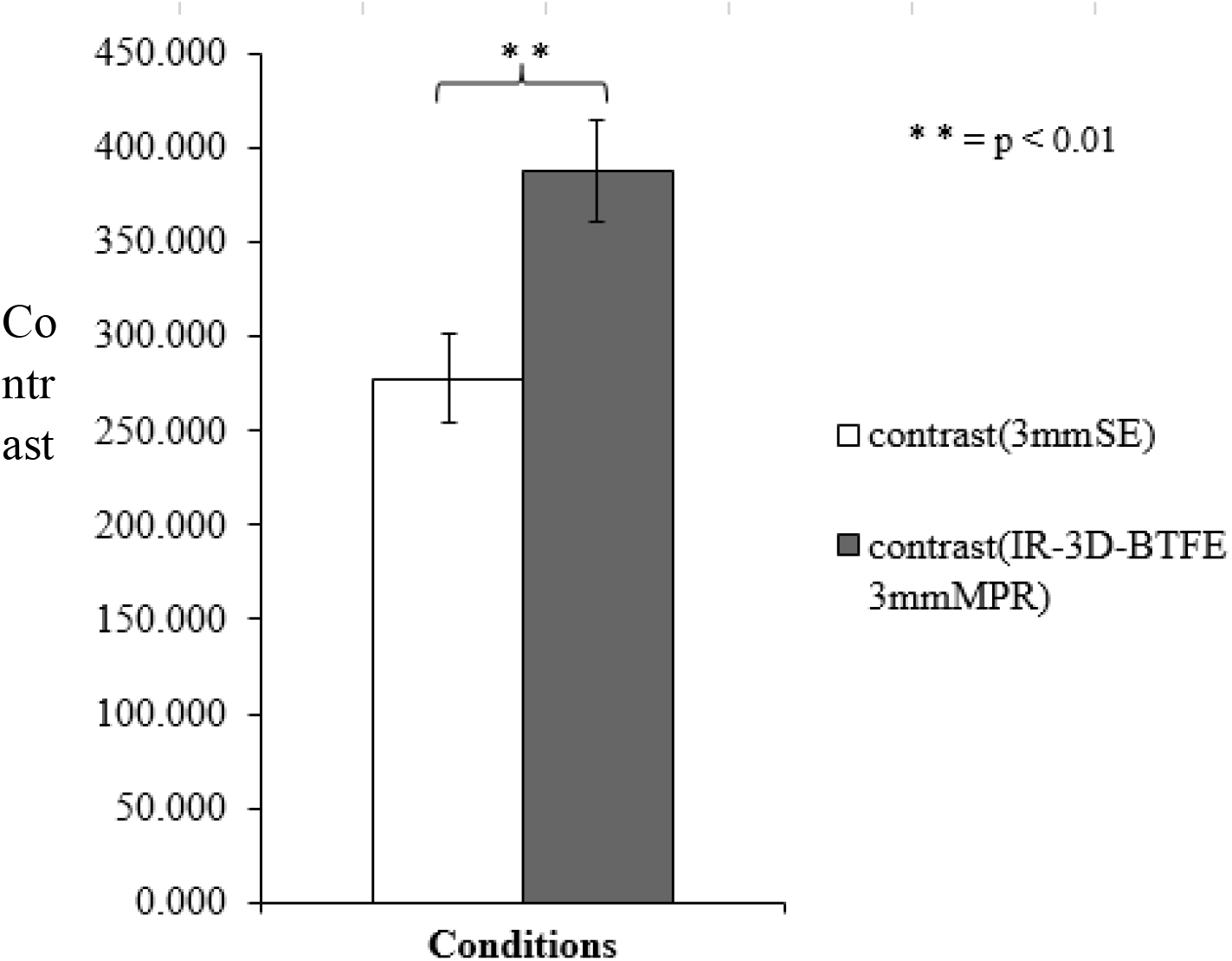
shows the difference between the average values of SE-T1WI and 3D-IR-BTFE (slice thickness 3 mm).

The difference of the average value of slice thickness 1 mm of SE-T1WI and 3D-IR-BTFE is shown in Fig. 4. The difference test of the mean value was t (19) = - 10.828, p <.01 (p = 0.0000000014), d = −1.656.

**Fig. 4:**
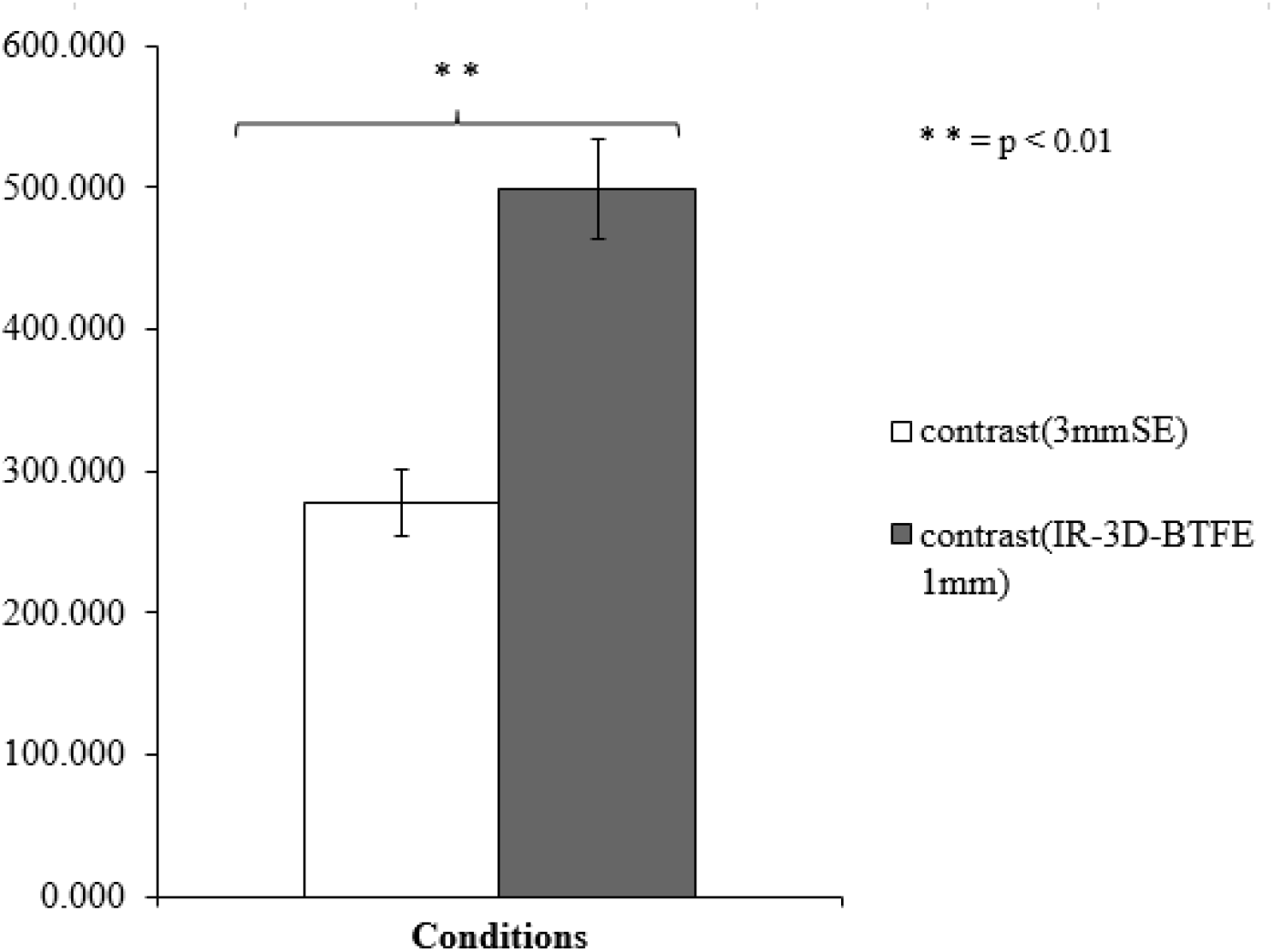
shows the differences between average values of SE-T1WI and 3D-IR-BTFE (slice thickness 1 mm).

The difference of the average value of 3D-IR-BTFE 3 mm and 3D-IR-BTFE 1 mm is shown in Fig. 5. The difference test of the average value was t (19) = −5.637, p <.01 (p = 0.000019), d = −787.

**FIG. 5:**
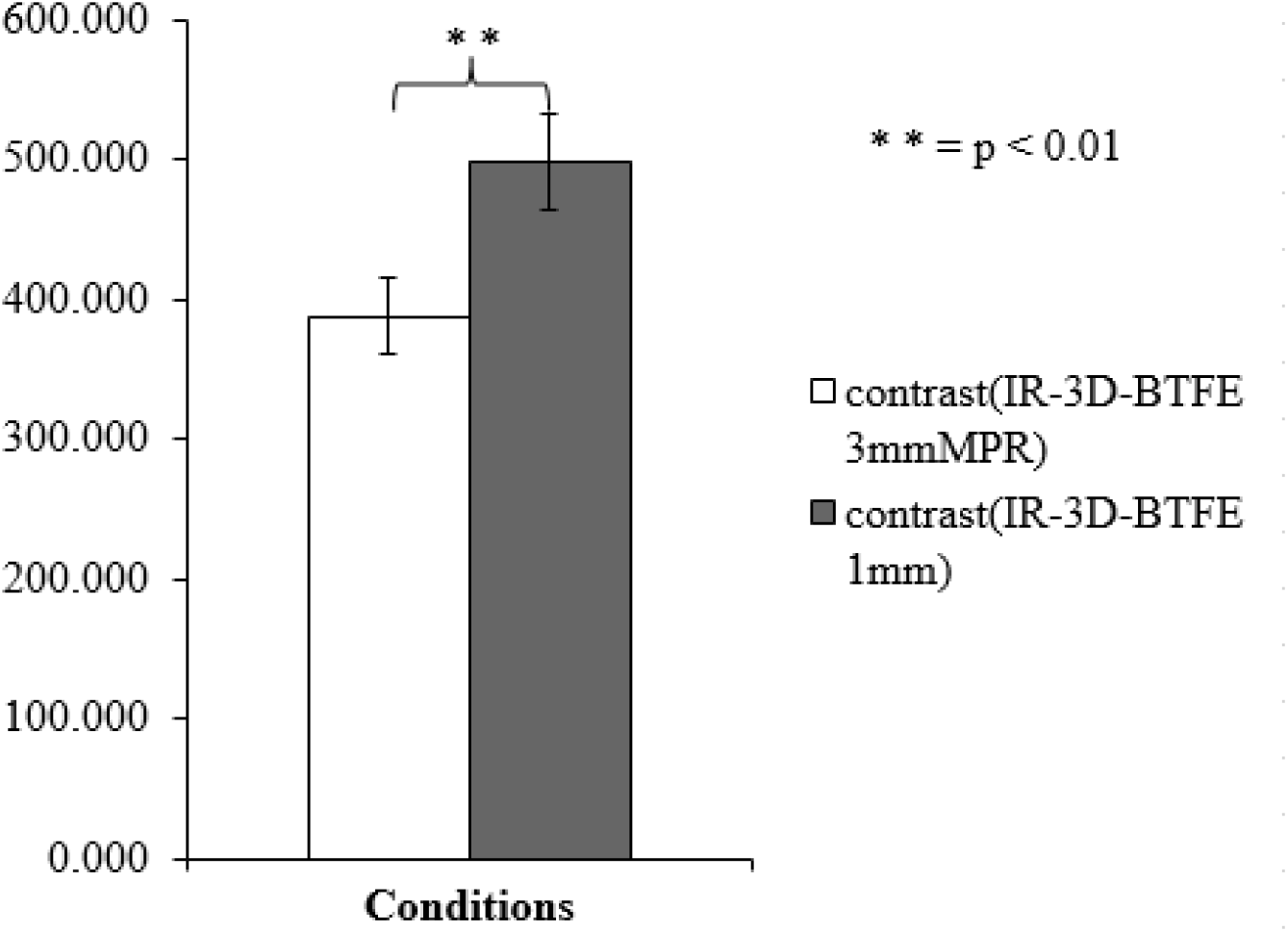
shows the difference between the average value of 3D-IR-BTFE (slice thickness 3 mm) and 3D-IR-BTFE (slice thickness 1 mm).

The difference between the slice thickness of 3 mm of SE-T1WI and IR-3 D-BTFE and the average of the ranks of IR-3 D-BTFE 1 mm (Friedman examination) is shown in Fig. 6. The difference test was z (14) = 36.100, p <.01 (p = 0.00000001), r = .602.

**FIG. 6:**
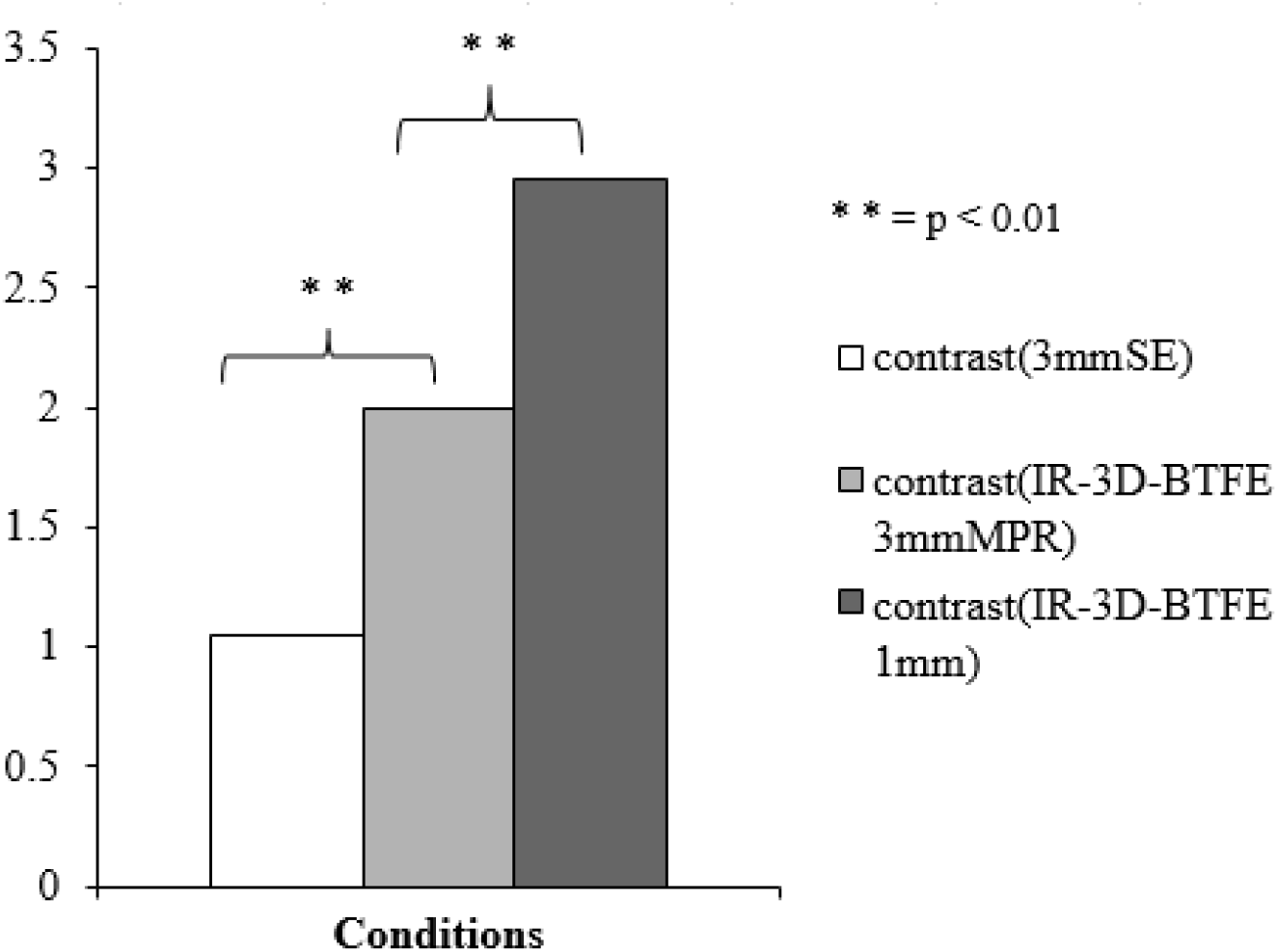
shows the results of Friedman’s test of the difference between SE-T1 WI and 3D-IR-BTFE (slice thickness 3 mm) and 3D-IR-BTFE (slice thickness 1 mm).

A contrast comparison of 3D-IR-BTFE and 3D-GRE is shown in Table 3. 3D-IR-BTFE has excellent contrast of 11 lesions per 12 lesions.

**Table 3:** shows a comparison of contrast between 3D-IR-BTFE and 3D-GRE.

|  | 3D-IR-BTFE |  |  | 3D-GRE |  |  | Contrast difference |
| --- | --- | --- | --- | --- | --- | --- | --- |
|  | Lesion | White matter | contrast | Lesion | White matter | contrast | 3D-IR-BTFEcontrast – 3D-GREcontrast |
| 1 | 2025.5 | 609.6 | 1415.9 | 1726.6 | 515.3 | 1211.3 | 204.6 |
| 2 | 1232.8 | 631 | 601.8 | 1113.2 | 531.7 | 581.5 | 20.3 |
| 3 | 1504.1 | 507.1 | 997 | 892.7 | 406.6 | 486.1 | 510.9 |
| 4 | 1196.6 | 507.1 | 689.5 | 819.3 | 406.6 | 412.7 | 276.8 |
| 5 | 1091.1 | 727.5 | 363.6 | 682.4 | 531.9 | 150.5 | 213.1 |
| 6 | 1320.8 | 891.8 | 429 | 887.1 | 558 | 329.1 | 99.9 |
| 7 | 1183.3 | 906.4 | 276.9 | 855.6 | 576 | 279.6 | -2.7 |
| 8 | 1254.1 | 793.3 | 460.8 | 712.5 | 587 | 125.5 | 335.3 |
| 9 | 1209.9 | 793.3 | 416.6 | 844.2 | 587 | 257.2 | 159.4 |
| 10 | 1919.4 | 770.9 | 1148.5 | 1189.6 | 476.9 | 712.7 | 435.8 |
| 11 | 1508.9 | 857.5 | 651.4 | 970.3 | 658.6 | 311.7 | 339.7 |
| 12 | 1329 | 626.1 | 702.9 | 1017.5 | 554.3 | 463.2 | 239.7 |

The difference in the average values of 3D-IR-BTFE 3 mm and 3D-GRE 3 mm is shown in Fig. 7. The difference test of the average value was t (11) = 5.247, p <.01 (p = 0.00027), d = 0.74. From this result, we can conclude that 3D-IR-BTFE has a better contrast than 3D-GRE.

**FIG. 7:**
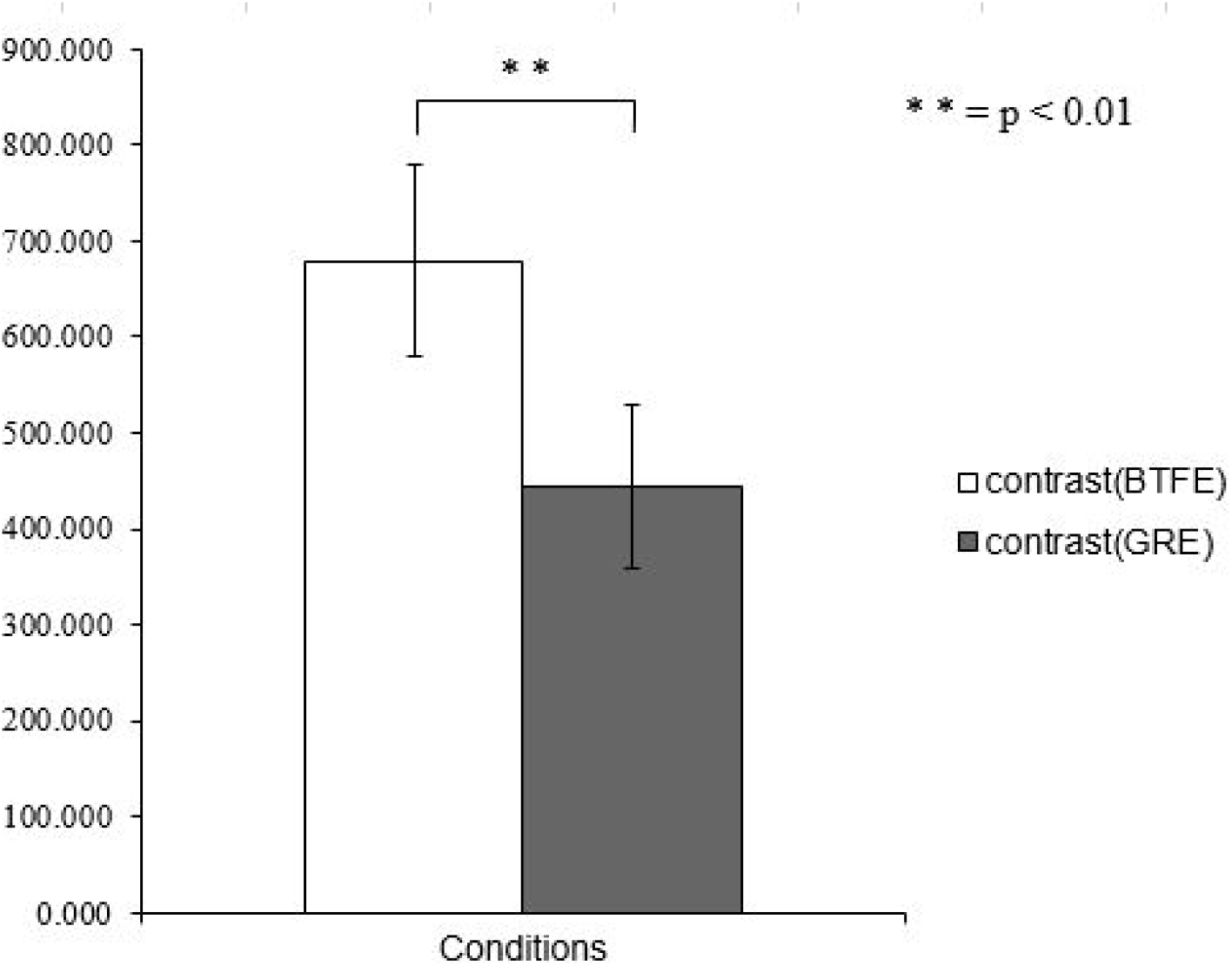
shows the difference between the average values of 3D-IR-BTFE and 3D-GRE.

## 5) Discussion

What is required in clinical diagnostic imaging of brain metastasis is to capture the presence of metastatic lesions, then the extent and size of metastatic lesions are important. If gadolinium contrast media is prescribed for the patient, I can reinforce contrast according to the T1 shortening effect. However, gadolinium contrast media and the T1 shortening effect are not necessarily in direct proportion. It is standard to prescribe 0.2 ml per 1 kg of the patient’s weight. For a brain metastasis lesion search, it is permitted to administer twice the amount of a specific gadolinium contrast agent only. The contrast enhancing effect of gadolinium contrast media has a limit. For this reason, we have to increase the contrast with the imaging method or static magnetic field strength. The factor that can raise the T1 contrast most in the imaging method is IR. 3D-IR-BTFE secured T1 contrast by adding IR.

In the image pick-up in two-dimensions, due to the characteristic of RF pulse, MRI equipment needs to prepare an interval between slices. This may be a challenge when searching for metastatic lesions- a challenge that could be solved if it could image in three dimensions. However, imaging in three dimensions increases imaging time. To eliminate this issue, a high signal-noise ratio imaging method must be selected. BTFE, in which TR is set short and imaged in a steady state without eliminating residual transverse magnetization, is an imaging method that can obtain excellent SNR and contrast. In the MRI apparatus studied here, the signal of the transition period can also be acquired. The signal of transition period can enlarge influence the T1 contrast. The T1 contrast can be increased by restoring longitudinal magnetization. To realize the longitudinal magnetization recovery, the parameter’s shot interval was lengthened to 4000 ms. I considered the possibility that the recovery of the longitudinal magnetization became larger by these, and excellent T1 emphasis was obtained. 3D-IR-BTFE considered SNR to be larger by BTFE than SE and gradient echo. It is likely that the good contrast was obtained by adding IR. To obtain further contrast, it is necessary to optimize IR and the numerical value of the shot interval.

Based on the above discussion, post-contrast 3D-IR-BTFE is superior to diagnose both brain metastatic lesions and the spread of metastatic lesions. It expected to be added to conventional SE-T1WI and 3D-GRE. From the restriction of SAR, the image pick-up is difficult with 3-Tesla equipment. In a high magnetic field device with 3-Tesla or more, it is necessary to investigate another imaging method that has IR added, and to a method of reducing the contrast medium.

## 6) Conclusion

This research was presented in September 2009 at the 37th Japanese Society of Magnetic Resonance in Medicine convention. Since the use of gadolinium contrast media has become increasingly strict in recent years, I am reporting this research. This study found that 3D-IR-BTFE with IR added was superior in contrast. It is possible to acquire images of thin slices in a shorter time than with the conventional imaging method and the Coherent-type gradient echo method could be used as a possible apparatus. To diagnose with the same degree of contrast as a spin echo or gradient echo, it is possible to reduce the contrast medium. The further examination is required for what contrast media loss in quantity is realized concretely. These results suggest that 3D-IR-BTFE, with excellent contrast, could be useful for other examinations at sites with little effect of motion.

